# Single-cell profiling resolves gain- and loss-of-function mechanisms in *CASR* to advance mechanism-aware variant classification

**DOI:** 10.64898/2026.09.26.754690

**Authors:** Samskruthi Reddy Padigepati, David Stafford, Yuya Kobayashi, Christopher A. Tan, Joseph Kim, Yunyun Jiang, Karen Ouyang, Jason Reuter, Alix M.B. Lacoste

## Abstract

Multiplexed assays of variant effect (MAVEs) and computational methods have advanced variant classification but have struggled to distinguish the molecular mechanism of variants with sufficient accuracy, limiting utility for multi-mechanism genes. Here, we coupled single-cell RNA-seq profiling with supervised machine learning to predict the mechanism of variants in *CASR*, a gene in which gain of function (GOF) variants cause hypocalcemia and loss of function (LOF) variants cause hypercalcemia. We engineered endogenous *CASR*-depleted HEK293 cells to express a single exogenous variant copy, measured their transcriptional profiles at single-cell resolution, trained a multi-class machine learning model on 96 expert-annotated variants (25 benign, 35 LOF, 36 GOF), and predicted mechanism for 157 variants of uncertain significance, of which 16 were predicted GOF and 54 LOF. The model showed high performance (macro F1 0.98, overall accuracy 0.98), and its predictions were corroborated by clinical phenotype and serum calcium measurements across more than 300,000 patients in a commercial genetic testing database. These results highlight the effectiveness of using molecular mechanism annotations with scRNA-seq to train models that distinguish molecular mechanisms of pathogenicity. The approach has immediate relevance for clinical variant classification and establishes a foundation for mechanism-guided clinical management in disorders with mechanism-dependent interventions, with potential application to other multi-mechanism genes.

## INTRODUCTION

Multiplexed assays of variant effect (MAVEs) have emerged as a powerful tool for clinical variant classification, enabling systematic functional characterization of thousands of variants in parallel.^1,2^ MAVEs quantify how genetic variants alter molecular and cellular phenotypes, including protein stability, enzymatic activity, and signaling, providing scalable experimental evidence to inform classification.^3^ When integrated with machine learning, MAVEs have resolved substantial fractions of variants of uncertain significance (VUS) in clinically important genes.^4–8^

Despite these advances, genes associated with multiple distinct diseases, each arising from different molecular mechanisms of pathogenicity, have not benefited to the same extent. In such genes, variants may cause disease through divergent functional effects—for example, loss-of-function (LOF) variants may underlie one disorder, whereas gain-of-function (GOF) variants cause another.^9^ Resolving the molecular mechanism is clinically significant, as accurate diagnosis, prognosis, and management often depend not only on whether a variant is pathogenic, but also on which disease it causes and through what molecular mechanism.

Capturing mechanism, however, is difficult both computationally and experimentally. Although *in silico* pathogenicity predictors have advanced remarkably, resolving the specific mechanism of a variant remains a substantially harder problem that current methods do not adequately address. Functional assays could help, but most MAVEs are gene-specific and centered on a single molecular readout, limiting their ability to distinguish variants that act through different mechanisms at scale. Recent advances, particularly single-cell RNA sequencing-based approaches, move beyond this constraint to assess variant effects through deep phenotypic profiling at single-cell resolution. ^10^ Because transcriptome-wide profiling provides a broadly applicable functional readout, these approaches have proven useful for supporting variant classification across genes with diverse molecular functions.^4^ Moreover, scRNA-seq–based assays have generated valuable functional evidence for genes in which either loss-of-function or gain-of-function mechanisms drive disease.^4^ Taken together, these findings suggest that scRNA-seq profiles could provide a general foundation for developing mechanism-specific variant predictions, particularly for genes associated with multiple diseases or distinct disease mechanisms.

Here, we present a generalizable framework for predicting disease-specific mechanisms of pathogenicity in genes associated with multiple disorders. We combine single-cell RNA sequencing-based variant effect measurements with supervised machine learning, leveraging expert-curated sets of variants with known disease associations. We selected the calcium-sensing receptor gene (*CASR*) as a model system because pathogenic variants in *CASR* are associated with two distinct clinical entities with opposing molecular etiologies: LOF variants cause familial hypocalciuric hypercalcemia type 1 (FHH1),^11^ characterized by mild hypercalcemia with inappropriately low urinary calcium excretion, while GOF variants cause autosomal dominant hypocalcemia type 1 (ADH1),^12^ marked by hypocalcemia with relative hypercalciuria. Using this system, we show that this integrated approach can not only identify pathogenic variants but also predict their specific disease mechanism, thereby enabling more precise clinical interpretation. This framework offers a path forward for resolving VUS in genes where multiple disease mechanisms complicate variant classification.

## MATERIAL AND METHODS

### Experimental approaches. Label curation

Variant pathogenicity and mechanism labels were curated by integrating three complementary annotation sources. First, variants classified as benign (B) in both a commercial genetic testing laboratory database and ClinVar were assigned a benign label, requiring concordance across the two sources to minimize the inclusion of misclassified variants. This yielded 889 benign variants. Second, mechanism annotations were compiled through expert review of internal and published evidence: 38 variants were annotated as LOF and 5 as GOF. Third, an additional 32 GOF annotations were incorporated from Roszko et al.,^13^ who systematically catalogued activating *CASR* mutations associated with autosomal dominant hypocalcemia.

### Variant selection

To construct a balanced training set for supervised learning, we selected a subset of labeled variants spanning all three mechanism classes: 26 benign, 38 LOF, and 37 GOF missense variants. For the LOF and GOF classes, we included all available mechanism-annotated variants. For the benign class, we randomly selected 26 missense variants from the larger pool. We included an additional 58 synonymous variants, not intended for model training, as an optional benchmark for quality control. 167 variants of unknown significance that are observed in multiple individuals and capable of being reclassified with high-quality functional evidence were selected to examine clinical actionability of model predictions.

### Derivation of the Recombination System

To introduce variants efficiently and precisely into cells, we chose a landing pad integration system that employs unidirectional BXB1 AttP/B recombination.^14^ We selected the T-REx™ 293 cell line as the parental background because it constitutively expresses the tetracycline repressor (TetR) and because we had prior success with related integration systems in this background^5^. We modified the T-REx™ 293 cell line (https://www.thermofisher.com/order/catalog/product/R78007) using a lentiviral transgene bearing an AttP site in-frame with Puromycin (https://www.systembio.com/pcdh-ef1-mcs-t2a-puro-cloning-and-expression-lentivector). Packaged landing pad transfer plasmid was transfected at a multiplicity of infection (MOI) greater than 1. Following two weeks of selection, we isolated and propagated clones. We individually assessed expanded clones for high integration efficiency and doxycycline-mediated transgene induction levels consistent with previous studies. We chose one optimal clone, henceforth referred to as the BXB1-AttP-293 parental cell line.

### Generation of the *CASR* variant library cell pool

The 326 selected variants were synthesized (Twist Bioscience) with variant-specific barcodes in the 3′ untranslated region of the transcript. Using the DNA HiFi Assembly Mix according to the manufacturer’s instructions (New England Biolabs) we cloned the barcoded variant pool into a donor plasmid backbone modified from pcDNA™5/FRT (https://www.thermofisher.com/order/catalog/product/V601020) in which the TetO sequence element (TCCCTATCAGTGATAGAGATCTCCCTATCAGTGATAGAGA) was positioned 3’ to CMV promoter and the FRT site was replaced with AttB (GGCTTGTCGACGACGGCGGTCTCCGTCGTCAGGATCAT). Analogous to the Flp-In™ system, BXB1-mediated recombination between the donor plasmid variant library (PVL) and the host landing pad AttP inserted a hygromycin resistance gene in frame. To maintain uniform distribution of variants, we plated rather than liquid-cultured the electroporated assembly (MegaX DH10B T1R Electrocomp™ Cells, Bio-Rad Gene-Pulser Xcell) and collected approximately 33,200 colonies for plasmid DNA isolation (Endo Free ZymoPURE II Plasmid Midiprep). To rule out synthesis and cloning errors and ensure high-quality starting libraries, we performed long-read sequencing of the PVL on a Revio system (Pacific Biosciences).

To generate the *CASR* variant engineered cell pool (ECP), we co-transfected the BXB1-AttP-293 parental cell line with a BXB1 expression plasmid (2:1 PVL:BXB1) using FuGENE 6 (3:1 FuGENE 6:total DNA). Forty-eight hours post-transfection, we cultured 5×10 cells with hygromycin (150 μg/mL) for 9 days to select for successful recombination. Based on colony counts and flow analysis for GFP on a non-selected parallel culture, we estimated the resulting ECP carried 500–800 unique integrations per variant (data not shown).

### Single cell capture, library prep, and sequencing

To prevent endogenous *CASR* expression from obscuring LOF variant effects, we depleted the native transcript 72 hours prior to single-cell capture by transfecting the ECP (Lipofectamine RNAiMAX) with a custom small interfering RNA (Horizon Biosciences) directed against the native transcript 3′ UTR (5’- GGAAAATGCTTCTGTTGTATT-3’), which is absent from the variant transgene. Forty-eight hours prior to capture, we applied 1 μg/mL doxycycline to induce variant expression. We confirmed both knockdown (76% reduction vs. parental cell line; data not shown) and induction (11.25-fold activation vs. non-induced ECP; data not shown) by quantitative PCR (Bio-Rad SsoFast™ EvaGreen® Supermix, Bio-Rad CFX Opus 96; primer sequences: *CASR* CDS FOR 5’-CCTGCATTGCCAAGGAGATC-3’; *CASR* CDS REV 5’- AAACACACCCAGCACAAAGG-3’; *CASR* UTR FOR 5’-TCTCCACGGTCAGATTTGCT-3’; *CASR* UTR REV 5’- ATGTCCCATCAGTCTGCACA-3’; ACTB reference 5’- CACCATTGGCAATGAGCGGTTC-3’ and 5’- AGGTCTTTGCGGATGTCCACGT-3’). We then performed single-cell capture following the manufacturer’s instructions (Chromium Next GEM Single Cell 3′ Kit, 10X Genomics), targeting 100 cells per variant. From the resulting cDNA, we generated both an expression sequencing library (manufacturer’s instructions) and a separate targeted variant–cell sequencing library to associate variant barcodes with cell barcodes (Figure S1).

### Computational analysis

#### Plasmid variant library quality control

We used long-read sequencing of the plasmid variant library to confirm associations between predetermined variant barcodes and their intended coding sequences with the variant of interest, and to exclude clones carrying synthesis or cloning errors. We extracted variant barcodes from long reads with cutadapt^15^ by matching 10-bp flanking sequences from the plasmid backbone (error rate 0.1) and retained reads with the expected barcode length of 8. We error-corrected extracted barcodes using the adjacency-graph clustering implementation in UMI-tools^16^ (Hamming distance: 2) and mapped them to the reference barcode library with UMI-tools whitelist mapping (Hamming distance ≤2).

We extracted coding sequence (CDS) inserts from the same reads with cutadapt using 20-bp flanking sequences (error rate 0.1) and retained inserts longer than 1 bp and shorter than 1.5 times the reference CDS length. We aligned each retained insert to the reference CDS using the striped Smith–Waterman algorithm implementation in scikit-bio^17^ and reported mismatches. For each barcode, we identified the most frequent mismatch (primary variant) and the second most frequent mismatch (secondary variant), and computed four QC metrics across supporting reads: total read count, primary variant ratio (fraction supporting the most common call), secondary variant ratio (fraction supporting the second most common call), and truncation rate (percentage of reads whose insert was >50 bp shorter than the median for that barcode). We retained a barcode when supported by ≥3 reads with a primary variant ratio >0.5, a secondary variant ratio ≤0.25, and a truncation rate ≤10%. We discarded barcodes failing these criteria.

#### Gene-expression quantification

Single-cell gene-expression libraries were prepared on the 10x Genomics Chromium platform and sequenced on an Illumina NovaSeq instrument. Raw sequencing data were processed with Cell Ranger version v8.0.0^18^ for demultiplexing, alignment to the human GRCh38 reference (refdata-gex-GRCh38-2024-A), and barcode and UMI counting, yielding a cell-by-gene UMI count matrix. The two sequencing captures were combined using the Cell Ranger aggregation pipeline with its built-in read-depth normalization.

#### Cell quality control

We identified low-quality cells using an unsupervised clustering approach on cell-level quality metrics. Prior to feature computation, we removed genes detected in fewer than 5% of cells and cells with zero total counts across the remaining genes. We then computed four features for each cell: total UMI count, number of genes with non-zero expression, mitochondrial UMI fraction, and total UMI count across the top 1,000 most highly expressed genes (ranked by their normalized expression sum across all cells). We log -transformed the total UMI and top-gene UMI features.

We scaled features with a robust scaler (median-centered, interquartile-range scaled) followed by min–max normalization to the [0, 1] range, and clustered them with HDBSCAN (hierarchical density-based spatial clustering of applications with noise).^19^ We selected HDBSCAN hyperparameters by grid search over min_cluster_size {10, 25, 50, 100, 200, 300, 400, 500, 600} and min_samples {10, 20, 30, 40}, choosing the combination that maximized HDBSCAN’s density-based clustering validation (DBCV) relative validity index. We classified clusters as low quality when ≥20% of cells had fewer than 5,000 total UMIs, and also flagged cells assigned to the HDBSCAN noise label (−1). We excluded all cells belonging to low-quality clusters from downstream analysis. This approach jointly captures cells with low RNA content, high mitochondrial fraction, and low gene complexity, which tend to co-cluster.

#### Variant-to-cell barcode association

To link each single cell to its expressed variant, we performed short-read sequencing on a dedicated variant–cell barcode (VCB) library in which each molecule contained both a 10x Genomics cell barcode and a variant barcode (Figure S1). We extracted cell barcodes (16 bp) and unique molecular identifiers (UMIs) from Read 1 (positions 0–15 and 16 onward, respectively), consistent with the 10x Genomics Chromium platform. We mapped observed cell barcodes to the Cell Ranger-validated cell barcode whitelist with UMI-tools (Hamming distance ≤2) and discarded unmapped barcodes. Within each cell, we collapsed UMIs with the UMI-tools adjacency algorithm (Hamming distance ≤1) to remove PCR duplicates.

We extracted variant barcodes from Read 2 with fuzzy regular-expression matching of the form (left_flank){error}([NACGT]{barcode_length})(right_flank){error}, using multiple 10-bp flanking patterns from the plasmid backbone to maximize sensitivity. We discarded reads producing conflicting variant barcode calls across patterns. We clustered extracted variant barcodes using adjacency-graph clustering implementation in UMI-tools and mapped them to the QC-validated variant barcode reference set within a Hamming distance of 1. To increase sensitivity, we additionally recovered reads containing variant barcode flanking sequences from the gene-expression library using the same extraction and error-correction procedure. We combined the reads from both the sources and deduplicated UMIs across the pooled set.

We assigned each cell to its primary variant by a UMI-ratio approach: for each cell, we counted the number of unique deduplicated UMIs supporting each variant barcode and nominated the variant with the highest ratio (UMI count for that variant divided by the total UMI count in the cell) as the primary variant call and second highest ratio as secondary variant call. We retained cells supported by ≥3 UMIs with a primary variant ratio >0.5 and a secondary variant ratio ≤0.25. The final output was a mapping of cell barcodes to variant identifiers, linking each single-cell transcriptomic profile to its introduced variant.

#### scRNA-seq data preprocessing

We integrated the UMI count matrix, cell quality-control labels, and cell-to-variant barcode assignments by retaining only high-quality cells with a successful variant assignment. We excluded synonymous variants and variants with computationally predicted splice effects (gain or loss of splice sites) called by Pangolin,^20^ since their transcriptomic profiles primarily reflect splicing disruption rather than the protein-level functional consequences of the intended missense or nonsense mutation. We excluded variants represented by fewer than five cells to ensure stable per-variant aggregate expression profiles for classification. We filtered genes using a per-variant adaptive threshold: we retained a gene if it was detected (non-zero UMI count) in more than 10% of cells in at least one variant group, preserving genes with biologically meaningful variant-specific expression even when expressed in only a subset of variants.

We processed the filtered count matrix with Scanpy.^21^ We normalized UMI counts in each cell to the median total UMI count across cells and log transformed the values. We identified the top 2,500 highly variable genes with the Seurat v3 method,^22^ selected within each sequencing capture and merged across captures. We subset the matrix to these genes, regressed out the linear effect of total UMI count per cell to remove residual library-size confounding, and scaled gene expression to zero mean and unit variance with values clipped at ±10 standard deviations.

We aggregated single-cell expression profiles to variant-level mean expression profiles by taking the arithmetic mean expression of each of the 2,500 highly variable genes across all cells expressing the same protein-level variant, yielding one 2,500-dimensional expression vector per variant.

#### Variant classification

We classified variants into three functional categories, benign (B), GOF, and LOF, using a nested leave-one-out cross-validation (LOOCV) framework with a principal component analysis (PCA)-support vector machine (SVM) pipeline implemented in scikit-learn.^23^ We used variants labeled B, GOF, or LOF (encoded as −1, 1, and 2) for training and nested leave-one-out cross-validated evaluation and predicted on unlabeled variants of uncertain significance.

In the outer loop, we held out each labeled variant once as the test sample, with all remaining labeled variants forming the training set. Within each outer fold, we optimized hyperparameters by 5-fold stratified cross-validation over a grid spanning PCA components, SVM regularization parameter C {0.1, 1, 10, 100, 1000}, kernel {linear, RBF}, and, for RBF kernels, gamma {“scale”, “auto”}. We set the number of PCA components adaptively to approximately 80% of the training set size, rounded down to the nearest multiple of ten, with candidate values spanning ±30 components in steps of ten (minimum of five), ensuring that the number of components never exceeded the number of training samples. We selected the best configuration by maximizing the one-vs-rest area under the receiver operating characteristic curve (AUROC).

For each outer fold, we retrained a fresh PCA-SVM pipeline on the full outer training set using the selected hyperparameters, with Platt scaling enabled in the SVM and no further probability calibration. We used the trained model to predict three-class probabilities for both the held-out labeled variant and all VUS. For labeled variants, we took the predicted class as the argmax of the probability vector from the single fold in which the variant was held out. For VUS, we averaged predicted probabilities from all outer folds and assigned the final class by argmax. We fixed all random seeds at 42 to ensure reproducibility.

#### Clinical Data Analysis

##### Study population

We identified patients carrying *CASR* variants through the Labcorp genetic testing database. The Western Independent Review Board protocol number CR-001-02 (Tracking ID 20161796) approved the use of de-identified patient data for all analyses.Three patient exclusion criteria were applied. First, patients carrying both a VUS included in this study and a Pathogenic/Likely Pathogenic (P/LP) variant in *CASR* — whether or not the P/LP variant was itself included in the study — were excluded to avoid confounding phenotypic attribution. Second, patients with a positive molecular diagnosis in any gene other than *CASR* were excluded to isolate the phenotypic effect of *CASR* variants. Third, to ensure each patient could be unambiguously assigned to a single variant classification category (Known GOF, Predicted GOF, Known LOF, Predicted LOF, Known Benign, or Predicted Benign), we resolved cases where a patient appeared in multiple variant categories as follows: GOF and LOF classifications were given precedence over benign classifications, such that a patient with assignments in a GOF or LOF category as well as a benign category was retained only in the GOF or LOF category. Any patient whose variants spanned more than one of the six classification categories after this reassignment — for example, carrying variants classified as both Known Benign and Predicted Benign — was excluded. For the serum calcium analysis, we matched the patients’ de-identified genetics results with their clinical laboratory dataset, including laboratory data from January 1, 2021 through June 1, 2026.

##### Clinical phenotype classification

We extracted clinician-provided ICD-10 codes and the indication for testing for each patient included in the study, and classified patients into four phenotype categories based on this information. The ADH1 category was defined by keywords including hypocalcemia and hypoparathyroidism, and ICD-10 codes and their descendants E20 (hypoparathyroidism) and E83.51 (hypocalcemia). The FHH1 category was defined by keywords including hypercalcemia and hyperparathyroidism, and ICD-10 codes and their descendants E21 (hyperparathyroidism), E83.52 (hypercalcemia), and R82.994 (hypercalciuria). To account for common typographical errors in clinical text (e.g., “hyperparathyrodism” for “hyperparathyroidism”), we used fuzzy string matching.

We followed a hierarchical approach for phenotype assignment. Cases were classified as ADH1 if ADH1 keywords were present without any FHH1 keywords, and as FHH1 if FHH1 keywords were present without any ADH1 keywords. If only one phenotype was indicated by ICD codes in the absence of keywords, cases were assigned to that phenotype. To resolve conflicts between keywords and ICD codes, we prioritized keywords, as the clinical indication text is considered more reliable. When both keyword sets were present, or when both ICD code sets were present, cases were classified as Both. All remaining cases were classified as Neither.

#### Serum calcium analysis

We extracted total serum calcium measurements (LOINC code 17861-6) for any patients with such a measurement in the linked genetics-laboratory data. For each patient, we compared analyses using different lookback windows - i.e., restricting laboratory results to tests performed within fixed time intervals (such as 1 or 2 years) prior to their earliest *CASR* genetic test date - as well as using no date-based filtering on laboratory test dates, and found only minor differences in the results. We therefore applied no date filter on laboratory results to capture the largest possible number of patients and variants.

We classified each calcium measurement relative to its laboratory-specific normal range, applying default ranges according to the patient’s age and sex when laboratory-specific reference ranges were unavailable.^24^ We then assigned each patient an overall calcium range status: in range (all measurements within normal limits), above normal (at least one measurement above normal, none below), below normal (at least one measurement below normal, none above), or conflicting (both above-normal and below-normal measurements present).

We grouped patients by variant category (Known Benign, Known GOF, Known LOF, Predicted Benign, Predicted GOF, and Predicted LOF) and computed the number and percentage of patients in each calcium range status category.

#### Statistical testing

To assess whether the distribution of clinical phenotype categories (ADH1, FHH1, both, or neither) and calcium measurement categories (above range, below range, in range, or conflicting) differed significantly across variant classification groups, we performed a permutation test for each pairwise combination of categories. For each pair, we computed a test statistic equal to the sum of absolute differences in mean percentage across all categories between the two variant groups. To ensure that variants with larger patient populations did not disproportionately influence the results, we first calculated the mean percentage for each individual variant (by averaging across all patients carrying that variant), then compared these variant-level means between groups, giving each variant equal weight in the analysis. We compared the observed test statistic against an empirical null distribution generated by 10,000 random permutations, in which variant labels were randomly reassigned between the two groups while preserving their respective sizes. This permutation approach yields exact p-values without relying on asymptotic approximations, which are unreliable when the number of variants per group is small. We adjusted p-values for multiple comparisons using the Benjamini– Hochberg false discovery rate procedure across all tested pairs.

#### Clinical impact

##### Variant classification

We analyzed germline DNA variants from individuals referred by clinicians for diagnostic genetic testing of hereditary disorders. We performed clinical variant classification, both before and after model training and deployment, using Sherloc, a validated points-based refinement of the ACMG/AMP guidelines.^25^

##### Reclassification assessment

To assess the impact of this work on *CASR* variant classification, variants that were classified as VUS at the time of model training were reinterpreted by Clinical Genomics scientists. The model was calibrated using positive and negative predictive values (PPV and NPV) and integrated into Sherloc in a manner consistent with our previous publication,^4^ in which 95% PPV (predicted GOF or LOF) and 95% NPV (predicted benign) corresponded to 2 pathogenic and 2 benign points, respectively, and 80% PPV or NPV corresponded to 1 pathogenic or benign point. VUS reclassification counts were tracked separately for each prediction category (GOF, LOF, and benign), and the number of patients who had previously received a VUS result as of November 13, 2025 was recorded.

## RESULTS

### Single-cell RNA-seq assay captures three distinct functional signatures for *CASR* variants

To systematically characterize the functional effects of select *CASR* variants, we performed a modified version of single-cell RNA sequencing (scRNAseq) based MAVE^4^ (Figure 1). We generated a barcoded variant library consisting of 326 variants in *CASR* that included benign, LOF, GOF and VUS *CASR* variants. We cloned the barcoded variant pool into a donor plasmid backbone and confirmed library quality by long-read sequencing. We then integrated the library into HEK293 cells using the BXB1 landing pad system and selected recombinants. Because HEK293 cells express *CASR*, we depleted endogenous expression using an siRNA directed against the endogenous *CASR* 3’ UTR that is absent from the variant transgene. We induced variant expression with doxycycline and performed scRNA-seq. In parallel, we sequenced a custom variant-cell barcode library for mapping variant identity to individual cells (see Methods).

**Figure 1:**
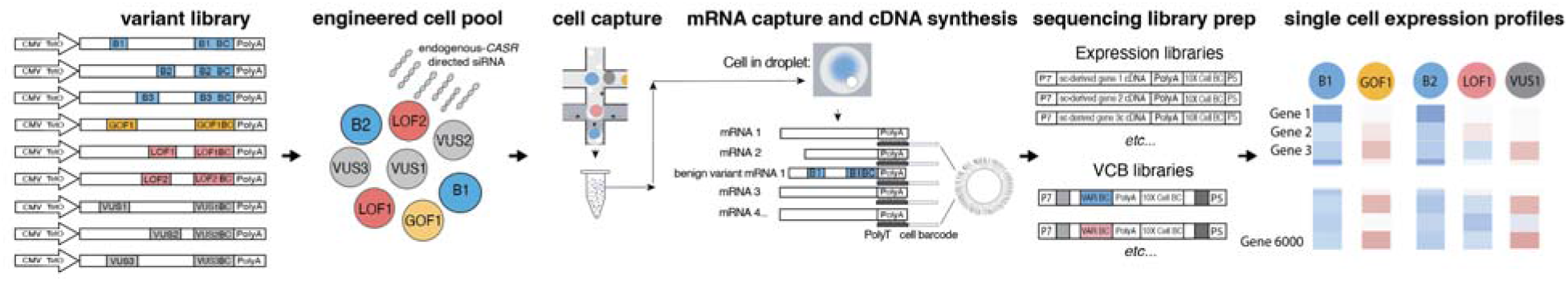
Single-cell RNA sequencing-based MAVE. A barcoded library of benign, LOF, GOF, and VUS *CASR* variants was integrated into HEK293 landing-pad cells. Following siRNA-mediated endogenous CASR knockdown and variant transgene induction, cells were encapsulated using the 10X Genomics GEM-X platform. Polyadenylated RNA was captured on bead-conjugated oligo-dT and reverse-transcribed, marking each cDNA molecule with a bead-specific barcode identifying its cell of origin. Gene-expression libraries measured the transcriptomic profile of each cell, while a separate variant-cell barcode (VCB) library was used to associate each profiled cell to its integrated variant (Figure S1). Expression profiles from variants with established mechanisms were used to train a multi-class machine learning classifier.

After cell quality control, assignment of cells to variants using the expression and enriched-barcode datasets and removing variants predicted to impact splicing (see Methods), we recovered 29,559 cells across 305 variants (67% of cells mapped to a variant barcode and passed QC). Excluding synonymous variants, 24,393 cells across 253 variants remained for downstream analysis (median 100 cells per variant, Figure S2). Of these 253 variants, 96 carried established clinical classifications (35 pathogenic LOF associated with familial hypocalciuric hypercalcemia Type 1 (FHH1), 36 pathogenic GOF associated with autosomal dominant hypocalcemia Type 1 (ADH1), and 25 benign); the remaining 157 were variants of uncertain significance (VUS). Lower-dimensional embeddings of the expression data, generated using uniform manifold approximation and projection (UMAP), revealed three distinct clusters corresponding to GOF, LOF, and benign variants (Figure 2A). We applied Leiden clustering on high-variance genes (see Methods), and the resulting heatmap revealed three coherent expression signatures, with benign, LOF, and GOF variants each sharing a distinct pattern of gene expression (Figure 2B).

**Figure 2:**
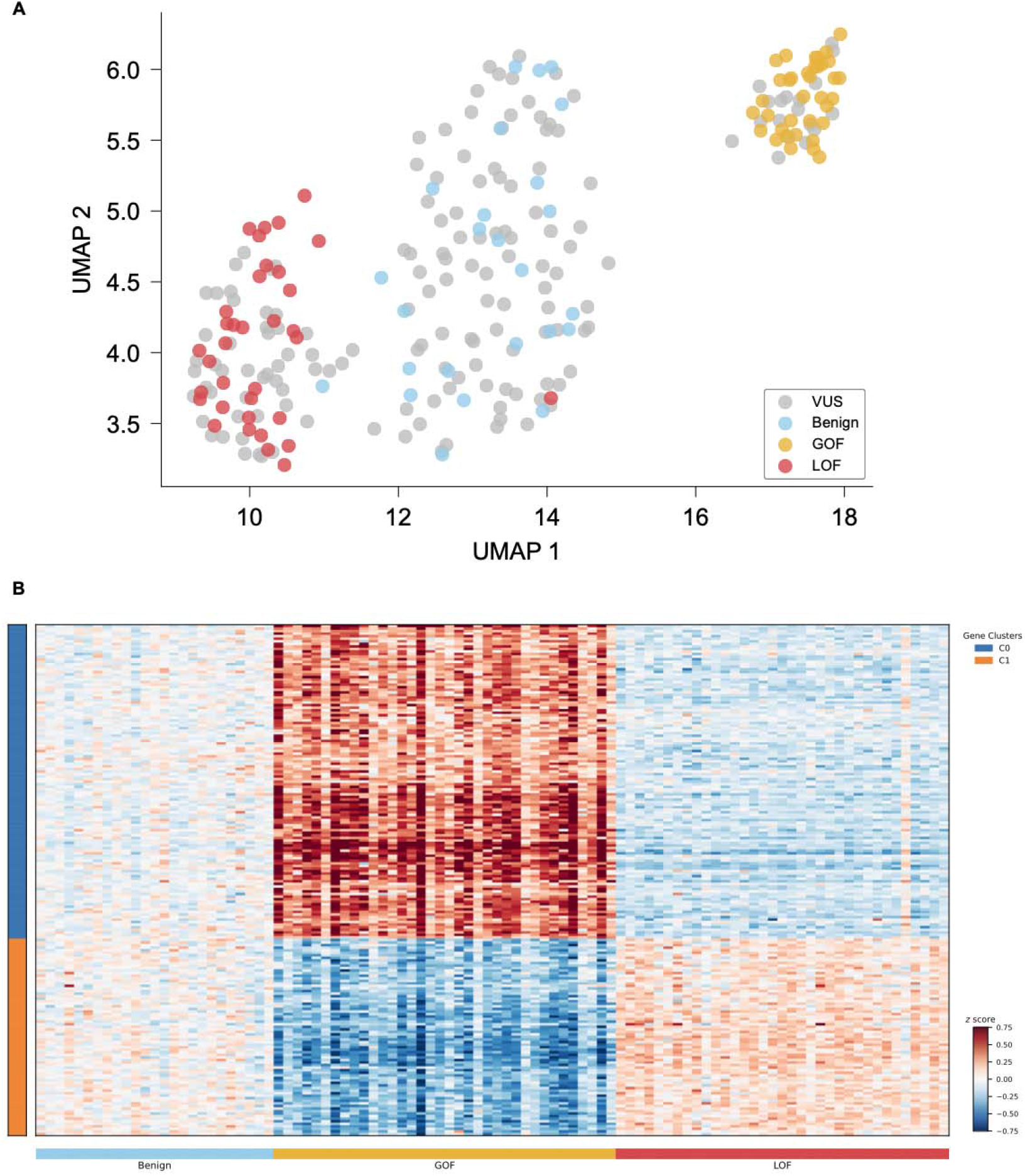
Expression signatures of *CASR* variants annotated by mechanism. (A) UMAP embedding of variant-level expression profiles (dots, n=253 variants), calculated by averaging expression of the top 2,500 highly variable genes across cells carrying each variant and colored by mechanism: light blue, benign missense; gold, GOF; red, LOF. Variants without an annotated mechanism (VUS) are shown in gray. (B) Mean scaled expression (z score, color bar) of the 200 most variable genes (rows) across the 96 mechanism-annotated variants (columns). Rows are grouped into two gene clusters (C0 and C1, left color bar); columns are ordered by mechanism (bottom bar) and codon position. Synonymous variants and VUS are not shown.

When we applied the same analyses to VUS, they partitioned into three clusters that closely mirrored the gene expression patterns of the characterized GOF, LOF, and benign variants (Figure 2A and Figure S3). The concordance between VUS clustering patterns and those of functionally characterized variants demonstrated that the VUS have interpretable functional consequences that can be classified using this approach.

### Machine learning (ML) model accurately predicts variant mechanism from functional data

To assess whether the scRNA-seq expression signatures could support variant classification, we trained a multiclass classifier to predict variant class (benign, LOF, or GOF) from variant-level mean expression profiles, using the 96 labeled variants (25 benign, 35 LOF, 36 GOF) as training data (see Methods, Figure 3A). To make full use of the limited labeled set while providing unbiased performance estimates, we evaluated the classifier using nested leave-one-out cross-validation, holding out each labeled variant in turn and predicting its mechanism from a model trained on all remaining variants.

**Figure 3:**
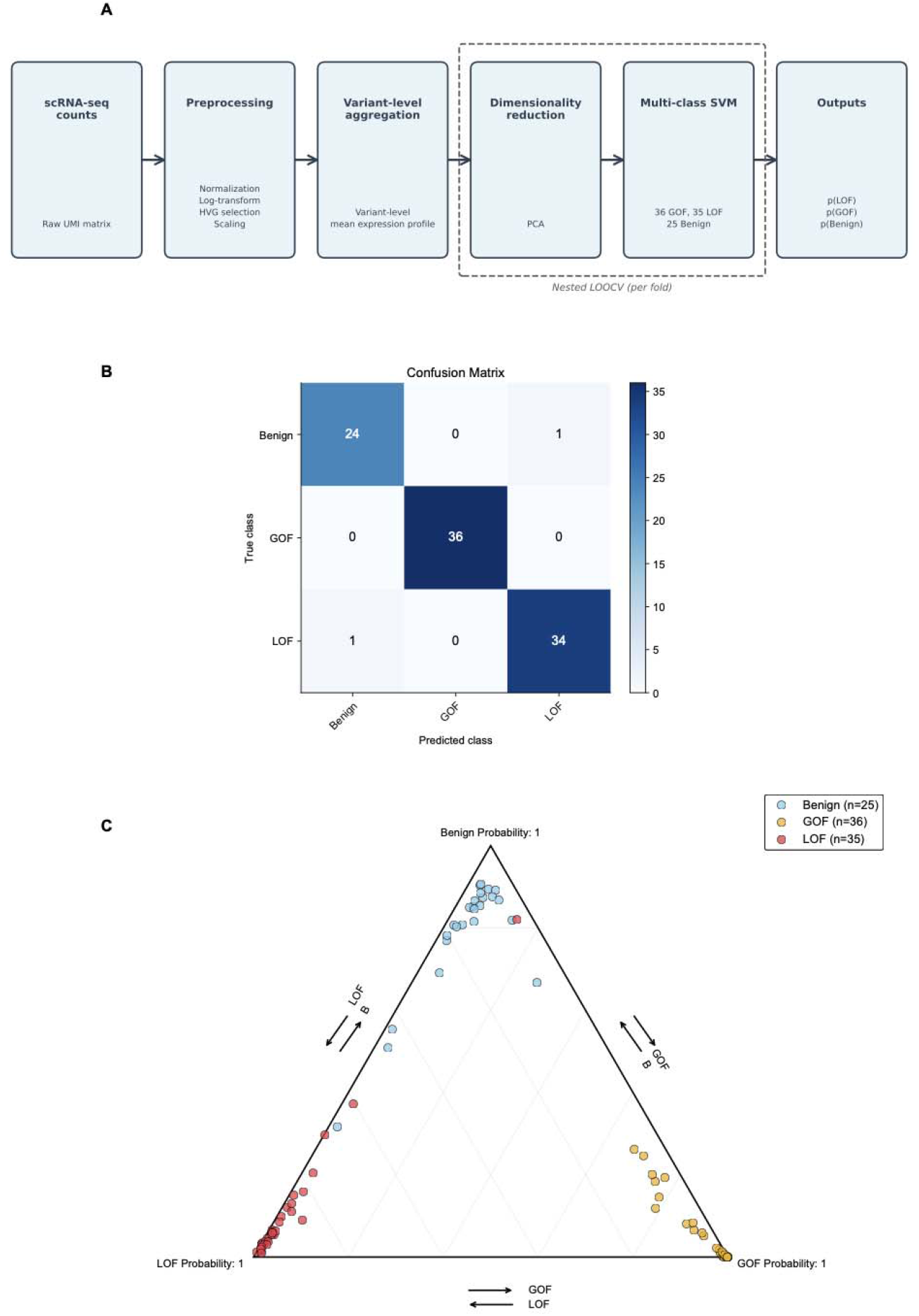
Prediction of *CASR* variant mechanism from expression profiles. (A) Schematic of the classification pipeline. Variant-level mean expression profiles were reduced by principal component analysis and classified by a multi-class support vector machine, with dimensionality reduction and classification nested within each leave-one-out cross-validation fold; outputs are per-class probabilities p(Benign), p(GOF), p(LOF). (B) Confusion matrix of nested leave-one-out cross-validation predictions for the 96 mechanism-annotated variants; overall accuracy 0.98. (C) Ternary plot of predicted class probabilities for n=96 variants (dots), positioned by p(Benign), p(GOF) and p(LOF), with proximity to a vertex indicating confident assignment: light blue, benign (n=25); red, LOF (n=35); gold, GOF (n=36). Arrows indicate the direction of increasing probability along each axis.

The resulting model recovered the known mechanism of held-out variants with high accuracy (overall accuracy 0.98), achieving per-class F1 scores of 0.96, 0.97, and 1.00 for benign, LOF, and GOF variants respectively, and one-vs-rest AUROC values ranging from 0.98 to 1.00 across the three classes (Figure 3B, Table S1, Figure S4, Figure S5). Misclassification was rare, limited to a single benign, and a single LOF variant exchanged across that boundary, while all GOF variants were correctly classified. When we visualized the predicted class probabilities for each variant, the labeled variants resolved cleanly toward their three respective vertices (Figure 3C, Figure S6). When applied to the 157 VUS, the model predicted 16 GOF, 54 LOF, and 87 benign (Table S2).

Each of the 253 variants is an SNV encoding a distinct amino acid substitution. Because different codons encoding the same amino acid substitution are likely to produce a protein with the same mechanism, we assigned each variant’s mechanism prediction to any other nucleotide changes encoding the same amino acid substitution. This expanded the set to 290 variants: 111 labeled (27 known benign, 42 known LOF, 42 known GOF) and 179 VUS (98 predicted benign, 62 predicted LOF, 19 predicted GOF), which we next evaluated against clinical evidence.

### Clinical and biochemical validation of predicted variant mechanisms

To validate that our model predictions correspond to clinically relevant disease mechanisms, we evaluated concordance between known and ML-predicted mechanism effects (benign, GOF, or LOF) by comparing their associated patient phenotype distributions. We performed two parallel analyses: first comparing clinical phenotypes extracted from clinical notes, and second comparing biochemical phenotypes derived from patient serum calcium measurements. To ensure an unbiased validation, we note that the model was trained exclusively on gene expression features and had no exposure to biochemical measurements or patient diagnoses.

To perform the clinical phenotype validation, we identified 300,095 patients from our database who carried *CASR* variants that were included in our functional assay and had documented clinical information, after excluding patients with both a VUS and a P/LP in *CASR* or a positive molecular diagnosis in another gene (see Methods). Phenotypes were classified using ICD-10 codes and text mining of clinician-provided indications, classifying patients as having ADH1, FHH1, both, or neither (see Methods).

Clinical phenotype correlated with known and predicted variant mechanism (Figure 4). In the predicted GOF variant category (n=83 patients, n=17 variants), the mean percentage of patients per variant showing a clinical phenotype consistent with ADH1 was 55.4 ± 11.0% (mean ± standard error), similar to the 54.9 ± 7.2% average observed for known GOF variants (n=76 patients, n=30 variants). Conversely, in the predicted LOF variant category (n=294 patients, n=41 variants), on average 53.3 ± 6.4% of patients had a clinical phenotype consistent with FHH1, compared to 59.9 ± 5.5% for known LOF variants (n=422 patients, n=34 variants). Finally, patients with predicted benign variants (n=1,254 patients, n=70 variants) showed an average of 96.05 ± 1.54% per variant having neither ADH1 nor FHH1 phenotype, similar to the 97.1 ± 0.6% average per variant for known benign variants (n=297,966 patients, n=25 variants) (Table S3).

**Figure 4:**
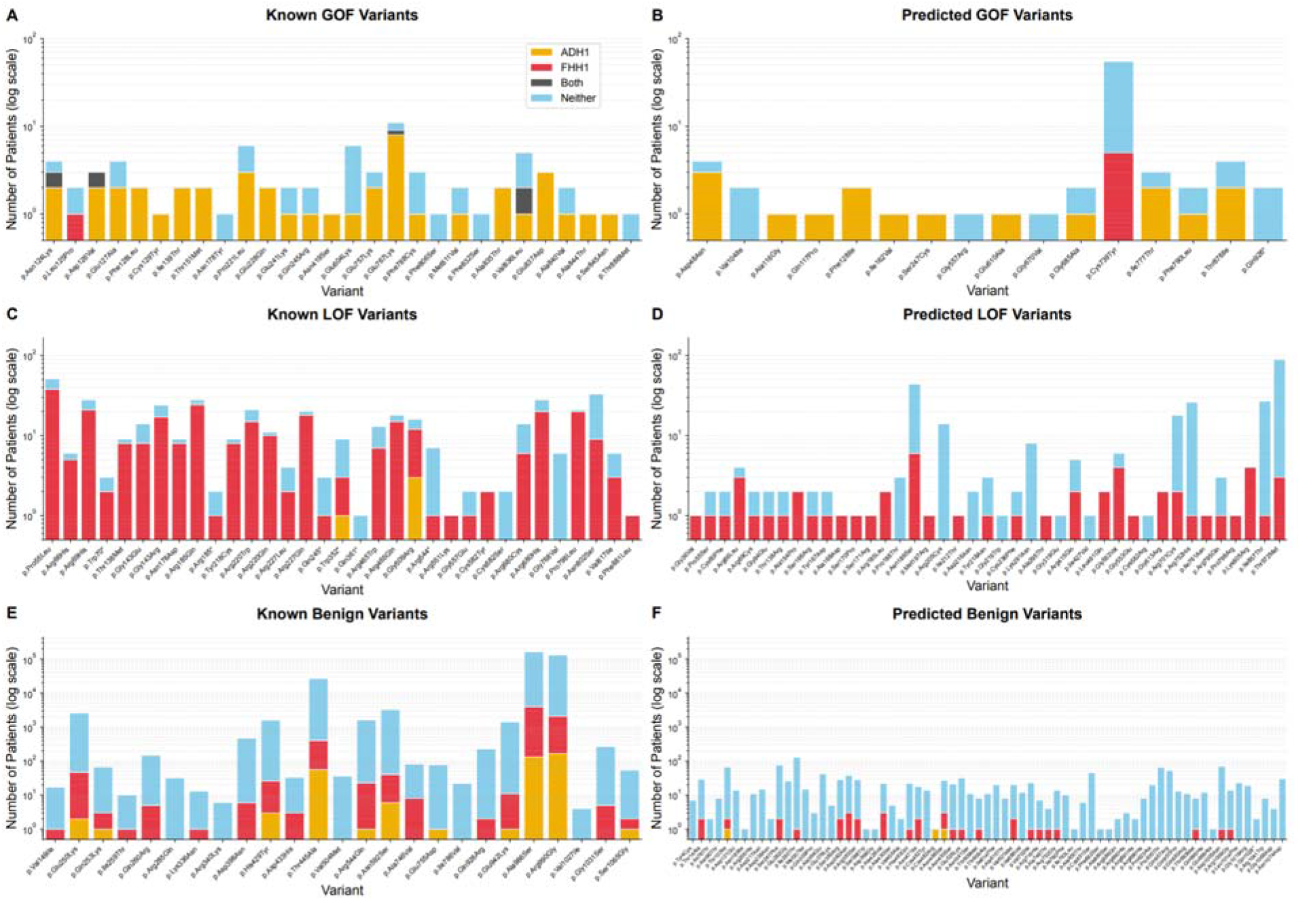
Clinical phenotype distribution across known and predicted mechanisms. Distribution of clinical phenotypes across *CASR* variants, stratified by variant classification and predicted mechanism. Stacked bar charts showing the number of patients with each clinical phenotype for *CASR* variants included in this study, in the log scale. Each bar represents a unique protein change variant grouped by predicted mechanism. Variants with the same protein change (i.e., same HGVS.p but different HGVS.c) were aggregated by summing patient counts. Colors indicate free-text indication fields and ICD10 codes consistent with the following clinical phenotypes: ADH1 (gold), FHH1 (red), Both ADH1 and FHH1 (gray), and Neither (blue). (A) Known GOF variants (n=30 unique nucleotide variants); (B) Predicted GOF (n=17); (C) Known LOF variants (n=34); (D) Predicted LOF (n=41); (E) Known benign (n=25); (F) Predicted benign (n=70). See also Table S3.

To provide biochemical validation, we analyzed serum calcium measurements across the patient cohort. This analysis included 78,243 patients after excluding patients with both a VUS and a P/LP in *CASR* or a positive molecular diagnosis in another gene, with 145 unique *CASR* variants. For patients with multiple calcium measurements, they were annotated as “in range” if all their measurements were in range, “above range” if at least one measurement was above range, and “below range” if at least one measurement was below range. If they had both above and below range measurements, they were classified as “conflicting” (see Methods).

Serum calcium levels correlated with known and predicted variant mechanism (Figure 5). Among patients with predicted GOF variants (n=26 patients, n=8 variants), the average percentage of patients per variant with at least one serum calcium level below normal range was 76.4 ± 15.5% (mean ± standard error), compared to 90.7 ± 6.3% for known GOF variants (n=14 patients, n=9 variants). Conversely, patients with predicted LOF variants (n=75 patients, n=22 variants) showed an average of 65.3 ± 9.6% per variant having above normal serum calcium, compared to 81.8 ± 6.1% average per variant for known LOF variants (n=114 patients, n=28 variants). Finally, patients with predicted benign variants (n=331 patients, n=54 variants) showed an average of 77.9 ± 3.5% per variant having serum calcium levels only in the normal range, compared to 84.1 ± 1.9% average per variant for known benign variants (n=77,683 patients, n=24 variants) (Table S4).

**Figure 5:**
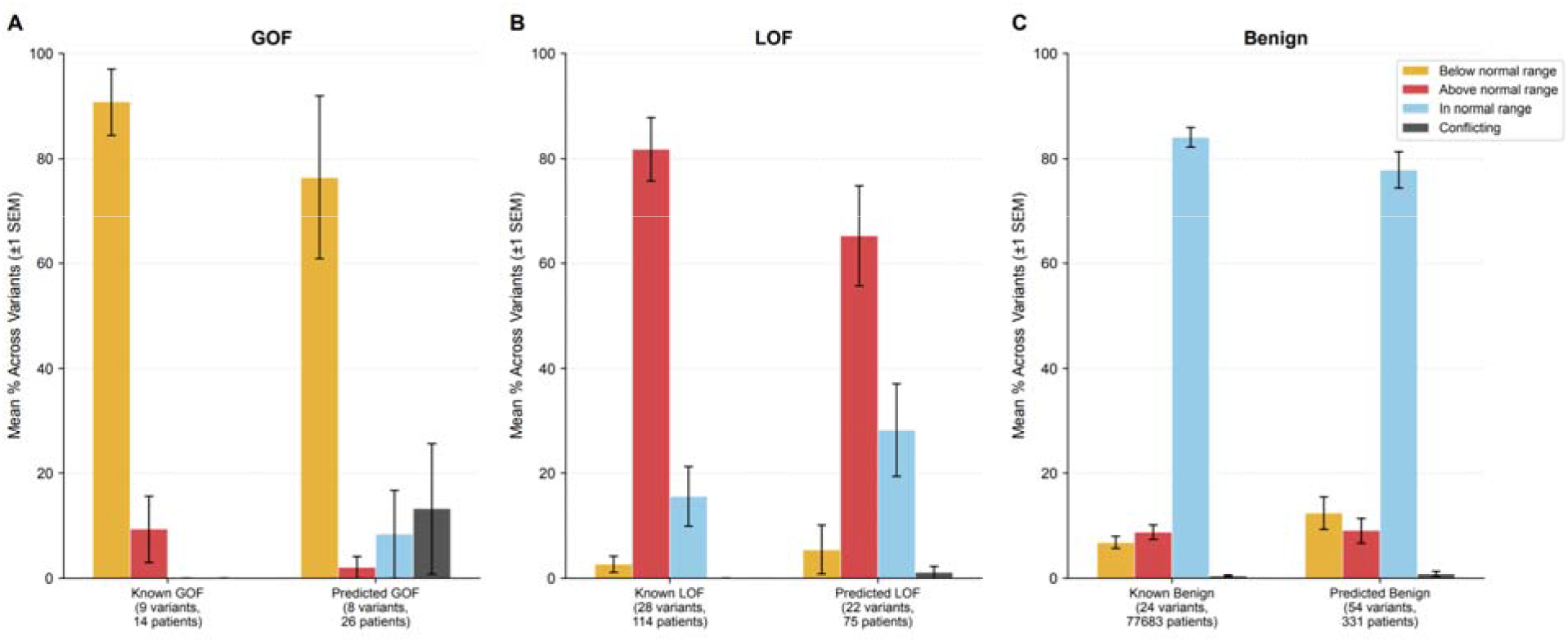
Distribution of serum calcium levels across *CASR* variant categories. Grouped bar chart showing the mean percentage of patients per variant with calcium measurements classified as below normal range (gold), above normal range (red), within normal range (blue), or conflicting (gray; patients with both high and low measurements). Error bars represent ±1 standard error across per-variant percentages within each category. Percentages are computed per variant then averaged across variants, giving equal weight to each variant regardless of patient count. 5A: Gain-of-Function variants (GOF label and predicted GOF mechanism), 5B: Loss-of-Function variants (LOF label and predicted LOF mechanism), 5C: Benign variants (Benign label and predicted Benign mechanism). Patient-level classification is based on all available calcium measurements. Numbers in parentheses indicate variant and patient counts per category. See also Table S4.

All nine cross-mechanism pairwise comparisons for both the phenotype and calcium analyses were significant at FDR < 0.01 with the Benjamini-Hochberg correction except for known GOF vs. predicted GOF (phenotype: p=0.96; calcium: p=0.36), known LOF vs. predicted LOF (phenotype: p=0.51; calcium: p=0.23), and known benign vs. predicted benign (phenotype: p=0.96; calcium: p=0.41).

Together, these clinical and biochemical validations demonstrate that functional predictions from scRNA-seq data are largely consistent with the *in vivo* consequences of *CASR* variants in patients. Nonetheless, phenotype consistency was stronger for the known vs the ML-predicted variant categories, suggesting lower penetrance of predicted variants, and highlighting the challenges of relying exclusively on clinical data for rare variant interpretation.

### Clinical impact: Large-scale VUS resolution affects thousands of patients

Having established the accuracy and clinical validity of the ML model, we evaluated the impact of incorporating it into clinical variant classification. We used the ACMG/AMP guidelines-based Sherloc framework to assign variant classifications (see Methods).

Prior to incorporating scRNA-seq data, the landscape of *CASR* variant interpretation included substantial uncertainty. Of the 290 variants in this study, 179 were classified as VUS using previously available evidence sources. We applied the functional predictions from our ML model to clinical variant classification following a performance-based calibration framework, as previously described.^4^ Briefly, model predictions were calibrated using positive predictive value (PPV) and negative predictive value (NPV) calculated against known pathogenic and benign variants (Figure S7, Table S2). Evidence strength was assigned based on PPV/NPV thresholds, and corresponding Sherloc evidence codes were applied to variants with prediction scores meeting performance criteria (see Methods). Variants were re-evaluated and classified within the context of all previously applied evidence.

Of the 179 VUS, 152 were observed in the Labcorp genetic testing database (87 predicted benign, 49 predicted LOF, and 16 predicted GOF). The addition of mechanism-specific functional data to these 152 VUS resulted in the reclassification of 61 variants (40.1%), resolving a substantial fraction of VUS (Figure 6A). Of the reclassified variants, 1 VUS was upgraded to likely pathogenic (LP) or pathogenic (P) based on GOF predictions consistent with ADH1, 24 VUS were upgraded to LP or P based on LOF predictions consistent with FHH1, and 36 VUS were downgraded to likely benign (LB) or benign based on neutral functional predictions (Figure 6B). Notably, 91 variants had mechanism-specific functional data applied and remained VUS. While the available evidence is currently insufficient, mechanism-specific functional data substantially increased the Sherloc score (Figure 6E), thus reducing the remaining burden of proof before their eventual reclassification.

**Figure 6.**
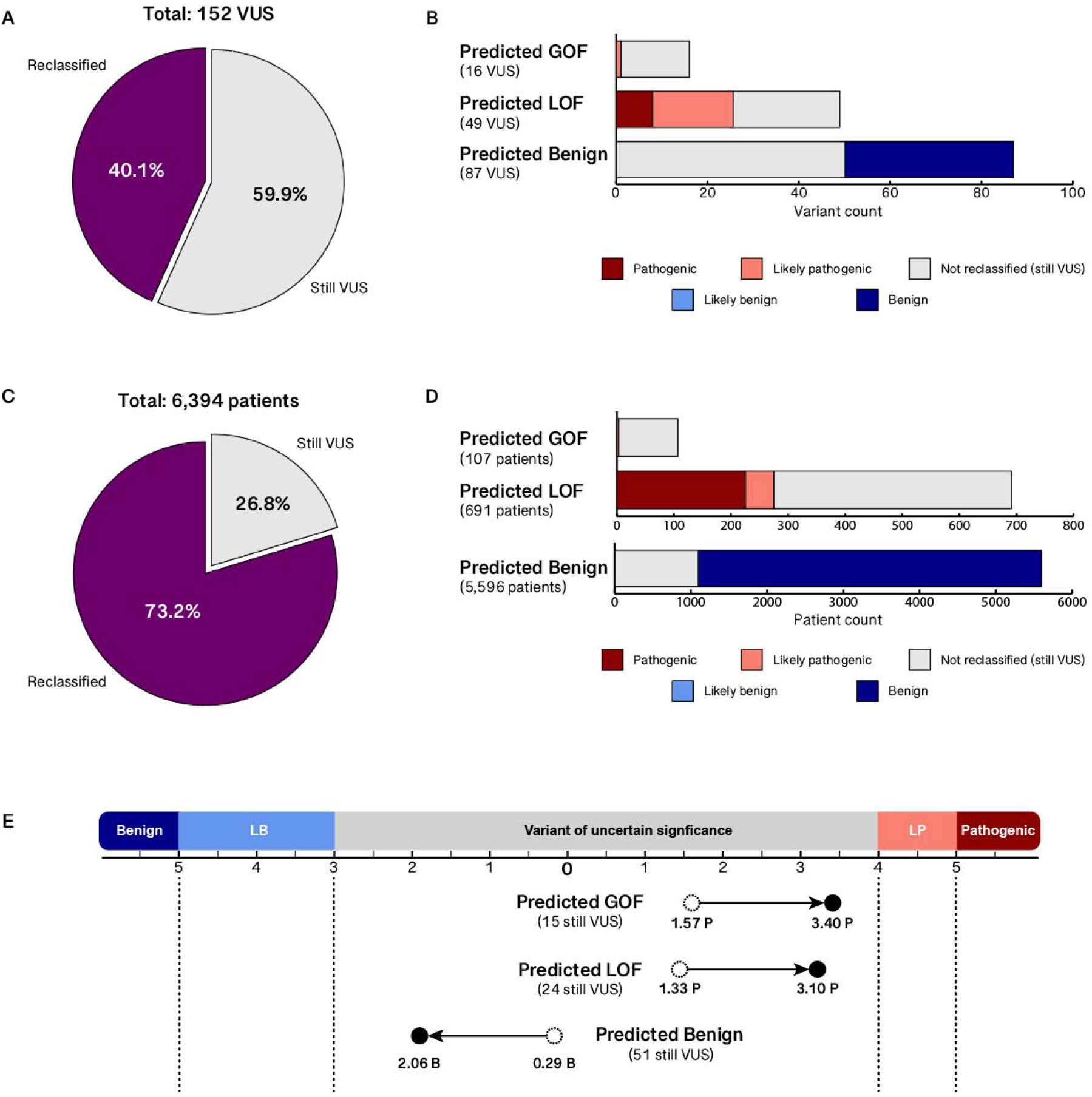
Impact of mechanism-specific functional predictions on *CASR* VUS classification and affected patients. (A) Pie chart depicts the proportion of *CASR* VUS reclassified to a non-VUS clinical classification following the incorporation of mechanism-specific functional evidence. (B) Stacked bar charts illustrate the distribution of reclassification outcomes, stratified by prediction type. (C) Pie chart depicts the proportion of patients carrying a *CASR* VUS whose variant was reclassified to a non-VUS clinical classification. (D) Stacked bar charts illustrate patient counts for each reclassification outcome, stratified by prediction type. (E) Mean Sherloc pathogenic scores (GOF and LOF predictions) and benign scores (B predictions), before and after incorporation of mechanism-specific functional evidence, for *CASR* VUS that remained VUS, stratified by prediction type.

To assess the real-world impact of the reclassified variants, we queried our database to identify all patients who had undergone clinical sequencing that included *CASR*. We identified 6,394 patients with *CASR* VUS detected through clinical testing. 73.2% of patients (4,679 individuals) carried at least one *CASR* VUS that was reclassified because of incorporating our scRNA-seq-based functional evidence into Sherloc (Figure 6C). Of these, 271 patients had variants reclassified from VUS to pathogenic or likely pathogenic, providing a molecular diagnosis where none existed previously (Figure 6D). An additional 4,408 patients had variants reclassified from VUS to benign or likely benign, decreasing diagnostic uncertainty (Figure 6D).

## DISCUSSION

Clinical variant classification is challenging for genes associated with distinct conditions that each arise from a different underlying molecular mechanism and require different clinical actions. For such genes, knowing whether a variant is pathogenic is not sufficient; the mechanism or associated condition must also be identified. Yet such annotations remain scarce. Clinical observation data are the gold standard for identifying the associated condition for a given variant. However, it is largely limited to variants with recurrent observations, making it unreliable for rare variants. While *in silico* predictors often have genome-wide coverage, contemporary predictors report only pathogenicity. Multiplexed functional assays offer a potentially scalable path forward but face a key limitation. Most MAVEs are designed around readouts associated with a single molecular function, so the multiple mechanisms by which variants act may not be captured. Moreover, because each assay is tailored to a specific gene, extending the approach across many genes is difficult.

To address these limitations, we developed a framework coupling scRNA-seq-based MAVE with a multi-class machine learning classifier trained on variants with curated mechanism annotations. Importantly, this approach is expected to be scalable across variants and genes, as a transcriptome-wide readout is capable of capturing multiple functional signals from a single experiment and is relatively gene agnostic. Explicitly modeling all known mechanism classes, rather than collapsing them into a binary pathogenic category, ensures each mechanism is learned as a distinct signal. High per-class performance on held-out variants then serves as evidence that each pathogenic mechanism has been accounted for, providing confidence in benign predictions. Applied to *CASR*, with GOF, LOF, and benign as the three classes, the classifier achieved F1 scores of 1.00, 0.97, and 0.96, respectively. Updating variants that were previously classified in the context of *CASR* genetic testing with model predictions resolved 61 of 152 VUS observed in the Labcorp genetic testing database, affecting 4,679 individuals carrying one of the reclassified variants.

Overall, model predictions were corroborated by both clinical phenotype and biochemical data, with mechanism-predicted variants showing distributions generally mirroring those of their known counterparts, providing independent validation that the model captures clinically meaningful functional distinctions. Predicted variants, however, showed weaker concordance with the expected phenotype compared to known variants on average. We interpret this generally as ascertainment bias rather than misclassification. Known LOF and GOF variants are known precisely because they segregated with disease in families and are therefore highly penetrant; the variants identified here likely went undescribed because their lower penetrance or milder biochemical effect kept them below the threshold of clinical ascertainment. In fact, a recent study that identified putative GOF variants found that these variants are associated with a milder form of ADH1.^26^ That such variants evade clinical detection even in a gene with phenotypically distinct presentations is precisely why functional evidence is a valuable complement to clinical data.

Our approach has two main limitations. First, readouts from cell-based assays do not always reflect a variant’s behavior in vivo. We, and others, functionally assessed p.Cys739Tyr to be GOF, but the available clinical and biochemical data do not align with the expected hypocalcemia.^27^ Crucially, incorporating functional evidence within a framework that also weighs clinical data prevented this signal from producing an erroneous classification, demonstrating the value of multi-evidence integration over reliance on any single data type. A second limitation stems from the supervised nature of our framework, which requires expert-curated variants spanning each mechanism class to train the classifier. For many genes, such labels are scarce or absent, restricting direct application of the supervised model. Our data suggest a path forward, the observed clear separation of *CASR* variants into distinct expression clusters without reliance on labels suggests that an unsupervised approach may be feasible: variants could first be grouped by transcriptional profile, with a small number of labeled examples used only to annotate the resulting clusters. This would extend mechanism-resolved profiling to genes where labeled variants are too few for supervised learning.

These limitations notwithstanding, extending this framework to more genes is the natural next step. This strategy holds particular promise for genes where clinical data alone struggles to distinguish the underlying mechanism. While *CASR* offers biochemically distinct phenotypes for each mechanism, some genes are associated with disorders that exhibit high phenotypic overlap despite different underlying molecular mechanisms. *SCN2A* is a prominent example: both GOF and LOF variants cause developmental and epileptic encephalopathy with seizures and intellectual disability, yet sodium channel blockers benefit GOF patients while worsening seizures in those with LOF variants.^28^ In such genes, where mechanism is not merely diagnostic but directs treatment, the need for mechanism-resolved functional evidence is greatest.

Expression-based functional profiling has previously been applied across many genes to predict pathogenicity^4^ and to distinguish mechanisms within a single gene.^10^ Our results show that combining mechanism-resolved functional profiling with supervised learning using expert-curated mechanism annotations can diminish diagnostic uncertainty at scale and enable actionable guidance for disorders in which the mechanism of pathogenicity determines the appropriate clinical response.

## Supporting information

Table S2. Model Predictions

## CONTRIBUTIONS

*Concept and Design:* Jason Reuter, Samskruthi Reddy Padigepati, David Stafford, Alix M.B. Lacoste

*Acquisition, analysis, and/or interpretation of data:* Samskruthi Reddy Padigepati, David Stafford, Jason Reuter, Alix M.B. Lacoste, Yuya Kobayashi, Joseph Kim, Yunyun Jiang, Karen Ouyang

*Administrative, technical and/or material support:* Samskruthi Reddy Padigepati, David Stafford, Yuya Kobayashi, Christopher A. Tan, Karen Ouyang, Jason Reuter, Alix M.B. Lacoste

*Supervision:* Jason Reuter, Alix M.B. Lacoste

*Writing – original draft:* Samskruthi Reddy Padigepati, David Stafford, Yuya Kobayashi, Alix M.B. Lacoste

*Writing – review and editing:* Samskruthi Reddy Padigepati, David Stafford, Yuya Kobayashi, Christopher A. Tan, Joseph Kim, Yunyun Jiang, Karen Ouyang, Jason Reuter, Alix M.B. Lacoste

## DATA AND CODE AVAILABILITY

The code to reproduce the analysis has not been deposited in a public repository because it is proprietary but can be made available from the corresponding author on request.

