## Supplemental Tables and Figures for "Single-cell profiling resolves gain- and loss-of-function mechanisms in *CASR* to advance mechanism-aware variant classification"

Table S1: Classification report

| **Class** | **Precision** | **Recall** | **F1-Score** | **Support** |
| --- | --- | --- | --- | --- |
| **Benign** | 0.960 | 0.960 | 0.960 | 25 |
| **GOF** | 1.000 | 1.000 | 1.000 | 36 |
| **LOF** | 0.971 | 0.971 | 0.971 | 35 |

Legend: Table S1: Classification report for CASR variant mechanism prediction. Per-class precision, recall, F1 score and support (number of annotated variants) from nested leave-one-out cross-validation of the PCA–SVM classifier across the 96 mechanism-annotated variants. Related to Figure 3B.

Table S3: Clinical phenotypes distribution by variant category

| **Type** | **Num Variants** | **Num Patients** | **% Patients with ADH1** | **% Patients with FHH1** | **% Patients with Both** | **% Patients with Neither** |
| --- | --- | --- | --- | --- | --- | --- |
| **Known GOF** | 30 | 76 | 54.9 ± 7.2 | 1.7 ± 1.7 | 3.4 ± 1.7 | 40.1 ± 7.2 |
| **Predicted GOF** | 17 | 83 | 55.4 ± 11.0 | 0.5 ± 0.5 | 0.0 ± 0.0 | 44.1 ± 10.8 |
| **Known LOF** | 34 | 422 | 0.9 ± 0.7 | 59.9 ± 5.5 | 0.1 ± 0.1 | 39.1 ± 5.4 |
| **Predicted LOF** | 41 | 294 | 0.0 ± 0.0 | 53.3 ± 6.4 | 0.0 ± 0.0 | 46.7 ± 6.4 |
| **Known benign** | 25 | 297966 | 0.2 ± 0.1 | 2.6 ± 0.6 | 0.0 ± 0.0 | 97.1 ± 0.6 |
| **Predicted Benign** | 70 | 1254 | 1.5 ± 1.4 | 2.5 ± 0.6 | 0.0 ± 0.0 | 96.0 ± 1.5 |

Legend: Table S3: Variant categories include known gain-of-function (GOF), loss-of-function (LOF), and benign variants based on established labels, and variants of uncertain significance (VUS) with machine learning-predicted mechanisms (GOF, LOF, or benign). Num variants: number of distinct nucleotide variants in each category with at least one patient with phenotype data. Num patients: total number of patients with phenotypic data after exclusions (see Methods). For each category, the remaining columns show the mean ± SEM percentage of patients in each clinical phenotype group, computed across individual variants: ADH1 (autosomal dominant hypocalcemia type 1), FHH1 (familial hypocalciuric hypercalcemia type 1), Both, or Neither, as determined by ICD-10 codes and text mining of clinician-provided test indications. Each variant is weighted equally regardless of patient count; ± values represent the standard error of the mean (SEM) across variants.

Table S4: Serum calcium distribution by variant category

| **Type** | **Num Variants** | **Num Patients** | **% Patients in range** | **% Patients above normal range** | **% Patients below normal range** | **% Patients with conflicting measurements** |
| --- | --- | --- | --- | --- | --- | --- |
| **Known GOF** | 9 | 14 | 0.0 ± 0.0 | 9.3 ± 6.3 | 90.7 ± 6.3 | 0.0 ± 0.0 |
| **Predicted GOF** | 8 | 26 | 8.3 ± 8.3 | 2.1 ± 2.1 | 76.4 ± 15.5 | 13.2 ± 12.4 |
| **Known LOF** | 28 | 114 | 15.5 ± 5.7 | 81.8 ± 6.1 | 2.7 ± 1.5 | 0.0 ± 0.0 |
| **Predicted LOF** | 22 | 75 | 28.2 ± 8.9 | 65.3 ± 9.6 | 5.4 ± 4.6 | 1.1 ± 1.1 |
| **Known benign** | 24 | 77683 | 84.1 ± 1.90 | 8.7 ± 1.4 | 6.8 ± 1.1 | 0.5 ± 0.2 |
| **Predicted Benign** | 54 | 331 | 77.9 ± 3.5 | 9.0 ± 2.3 | 12.3 ± 3.1 | 0.8 ± 0.5 |

Legend: Table S4: Variant categories are as defined in Table S3. Num variants: number of distinct nucleotide variants with at least one patient with calcium data. Num patients: number of patients with at least one serum calcium measurement after exclusions. For each category, the remaining columns show the mean ± SEM percentage of patients in each serum calcium status group, computed across individual variants. Calcium measurements (LOINC 17861-6) were classified relative to laboratory-specific reference ranges, applying default ranges according to the patient’s age and sex when laboratory-specific reference ranges were unavailable. Each patient was assigned a single calcium status: in range (all measurements within normal limits), above normal (at least one above-normal measurement, none below), below normal (at least one below-normal measurement, none above), or conflicting (both above- and below-normal measurements present). Each variant is weighted equally regardless of patient count; ± values represent the SEM across variants.

Figure S1

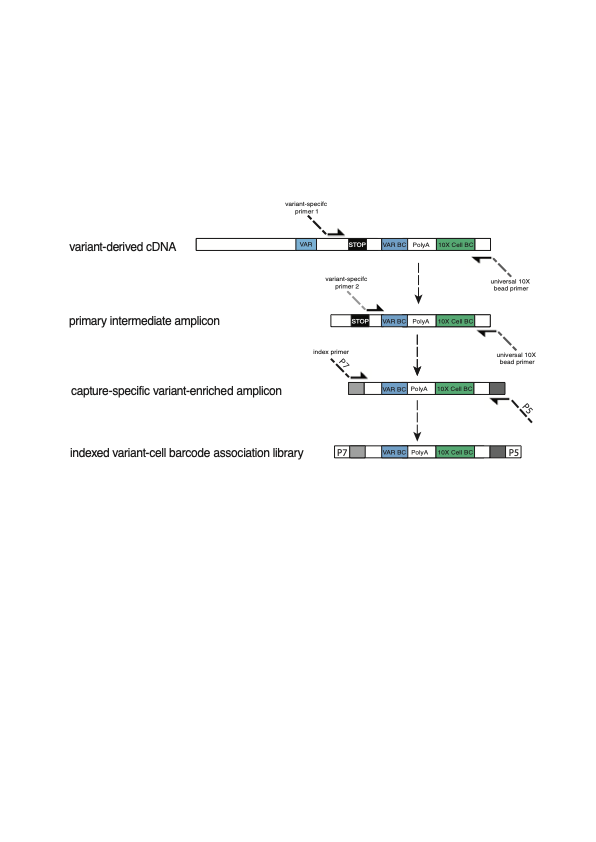

Legend: Figure S1: Custom variant-cell barcode library strategy.

Unique 10X genomics bead-specific barcodes are used to associate reads to cells. Within these single-cell transcriptomic profiles are cDNAs that derive from the exogenous variant. Because the expression library targets the 3' end of cDNAs, it is not possible to identify variants by direct examination of the coding sequence. As such, variants are programed with a variant-specific 8-bp barcode (Fig1A). However, due to the sparse matrix intrinsic to scRNA-seq, relying solely on variant-barcodes identified in the expression data results in missed cells. To address this feature of the workflow, we sequence a custom variant-cell barcode association library. Using pooled cDNA from a given capture as a template, an intermediate product is amplified using a primer set consisting of a target-specific forward primer and a universal 10X bead primer. This product is further refined with a second nested forward primer and the same 10X bead primer. Following bead-size selection, the second amplicon is indexed using a primer mix consisting of a custom SI-GA-E series P7 primer from 10X Genomics and a custom P5 primer.

Figure S2

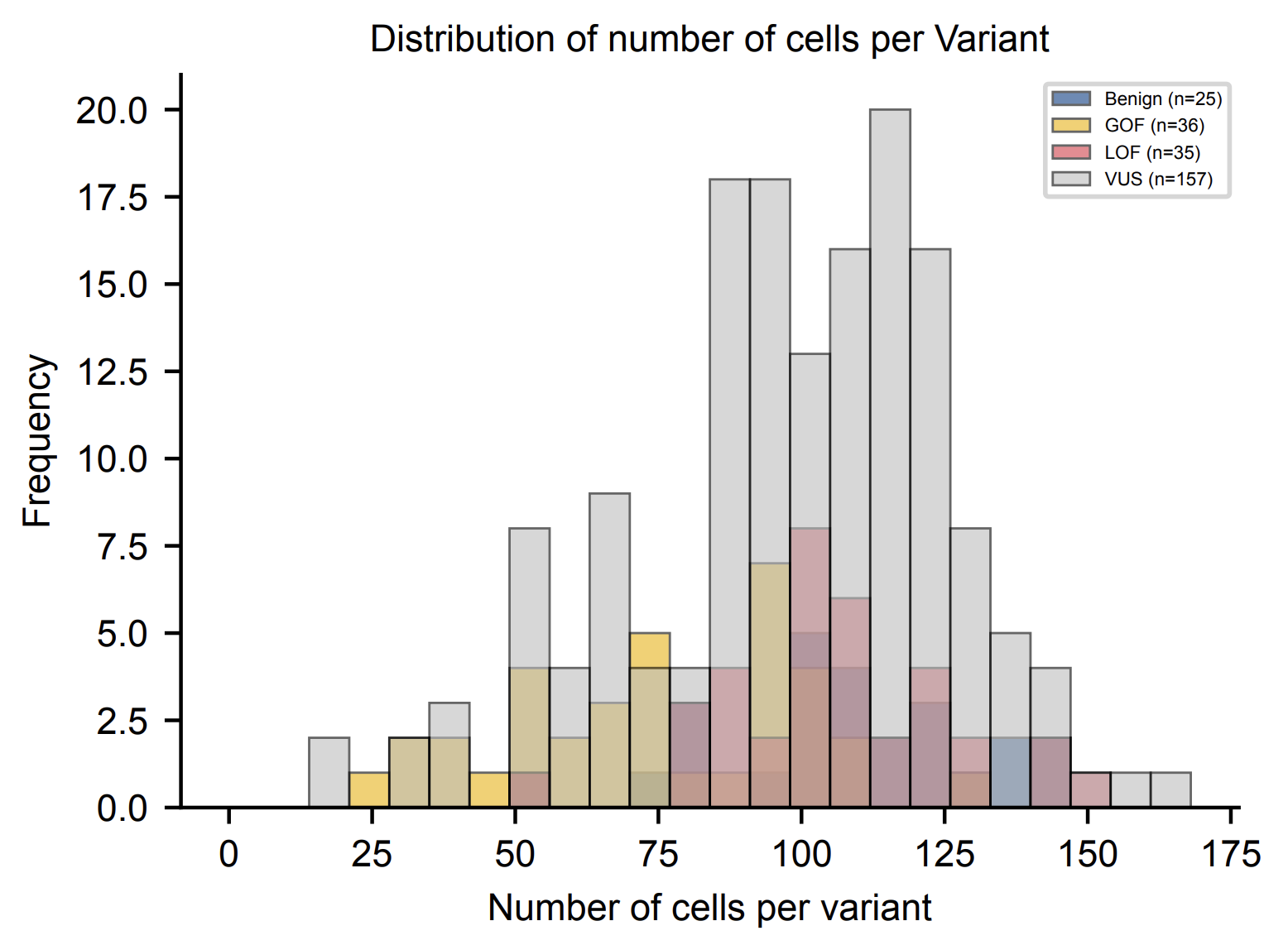

Legend: Figure S2: Distribution of cells per variant. Number of cells assigned to each variant after quality control and variant–cell barcode assignment, shown as overlapping histograms by mechanism: dark blue, benign (n=25); gold, GOF (n=36); red, LOF (n=35); gray, VUS (n=157). Median 100 cells per variant.

Figure S3

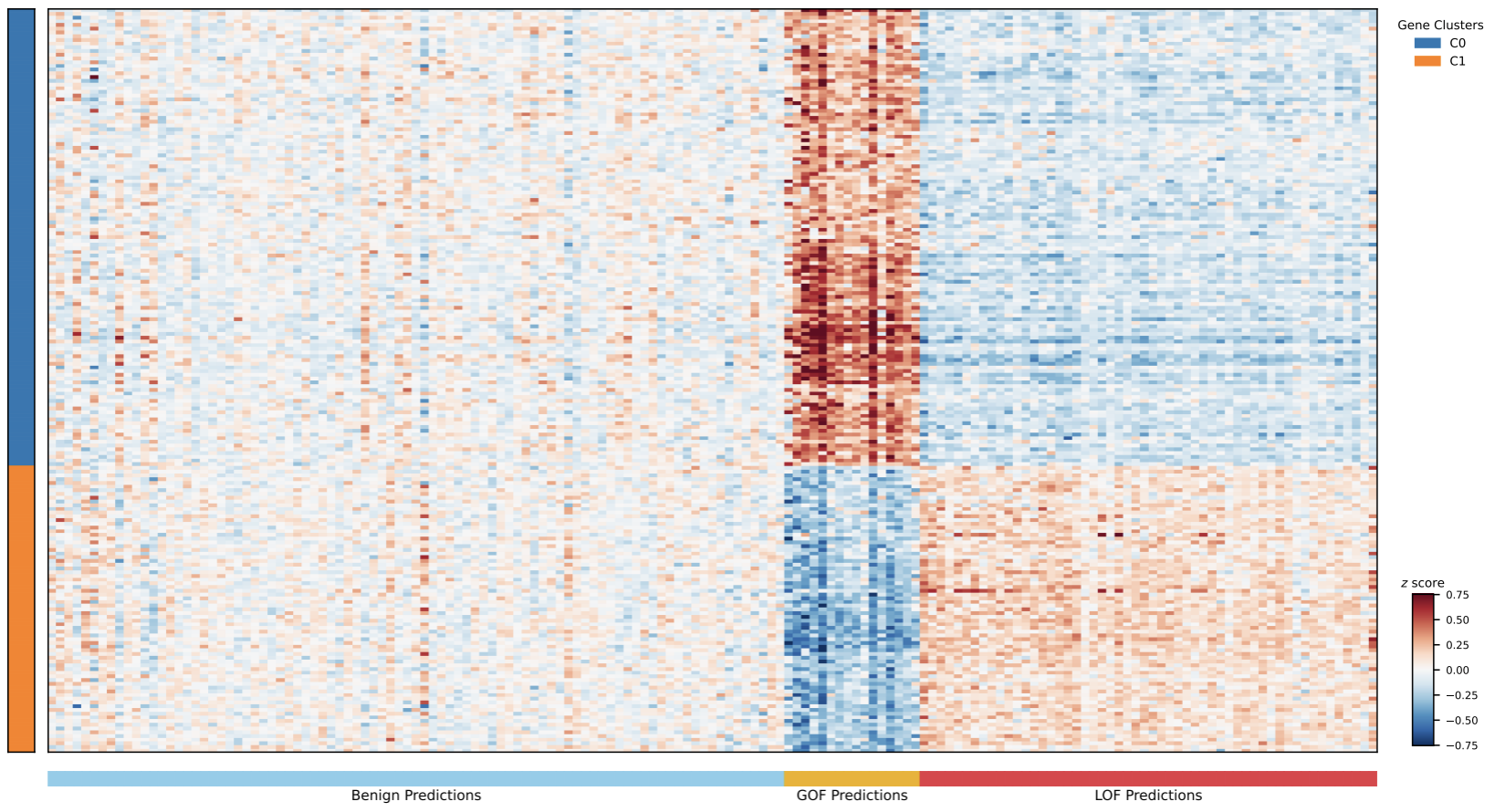

Legend: Figure S3: Expression signatures of CASR VUS grouped by predicted mechanism. Mean scaled expression (z score, color bar) of the 200 most variable genes (rows) across VUS (columns, n=157), ordered as in Figure 2B. Rows are grouped into two gene clusters (C0 and C1, left color bar); columns are ordered by predicted mechanism (bottom bar). VUS predicted benign, GOF and LOF show expression signatures resembling the corresponding annotated variants in Figure 2B.

Figure S4

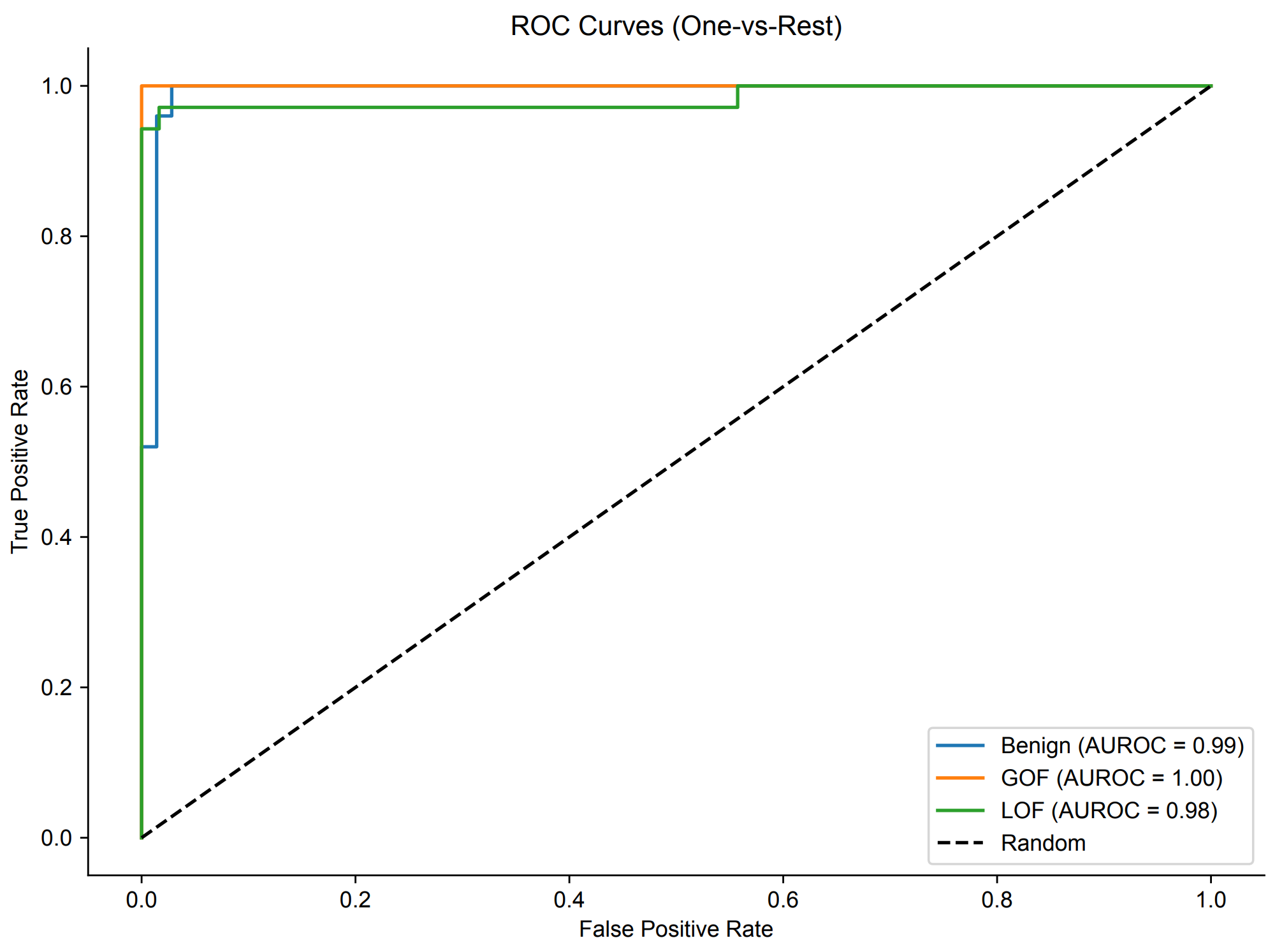

Legend: Figure S4: Classifier performance by mechanism class. One-versus-rest receiver operating characteristic curves from nested leave-one-out cross-validation of the PCA–SVM classifier across the 96 mechanism-annotated variants: blue, benign (AUROC = 0.99); orange, GOF (AUROC = 1.00); green, LOF (AUROC = 0.99). Dashed line, random classifier. Related to Figure 3B.

Figure S5

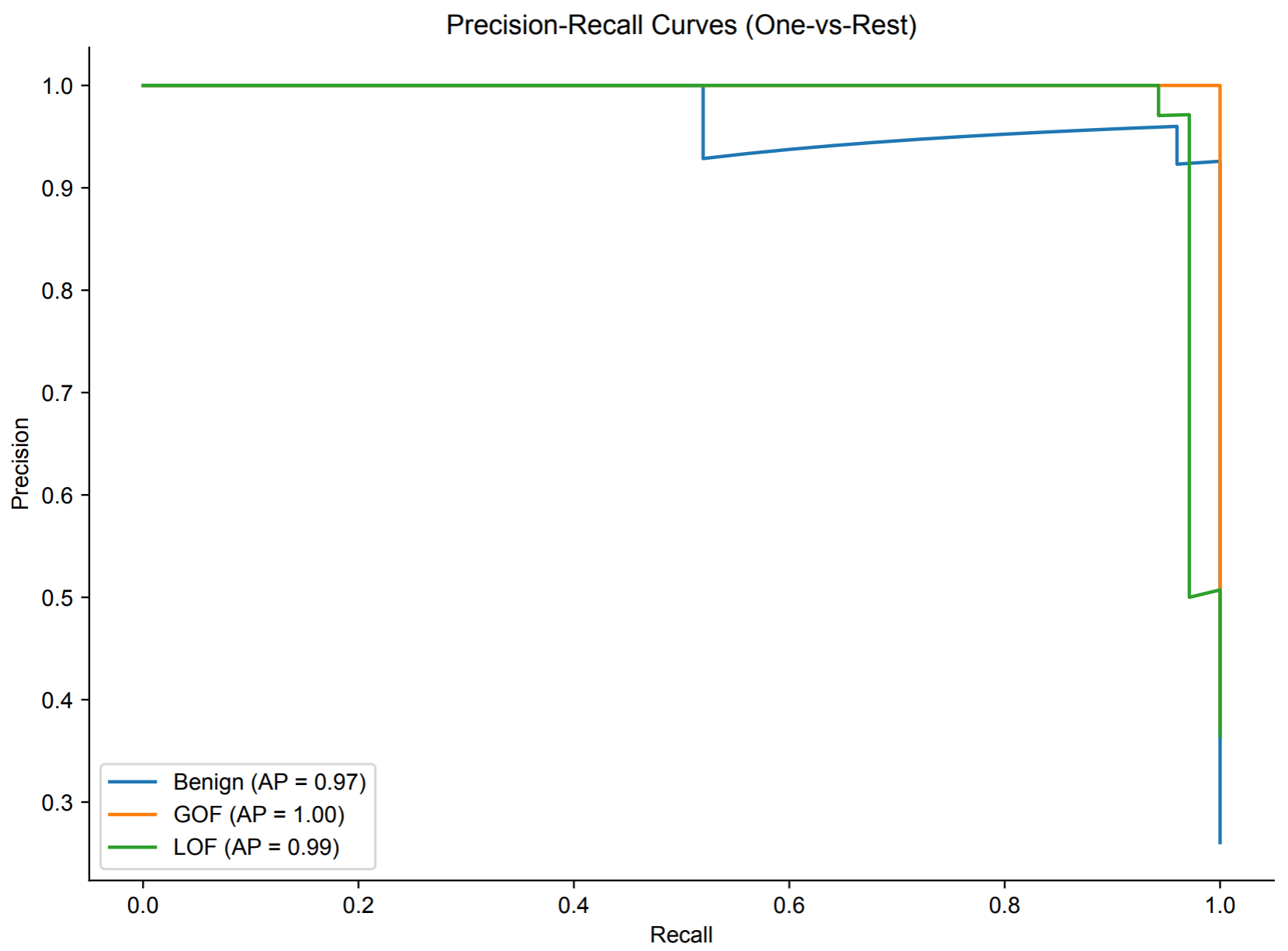

Legend: Figure S5: Precision–recall performance by mechanism class. One-versus-rest precision–recall curves from nested leave-one-out cross-validation of the PCA–SVM classifier across the 96 mechanism-annotated variants: blue, benign (AP = 0.98); orange, GOF (AP = 1.00); green, LOF (AP = 0.99). AP, average precision. Related to Figure 3B and Figure S4.

Figure S6

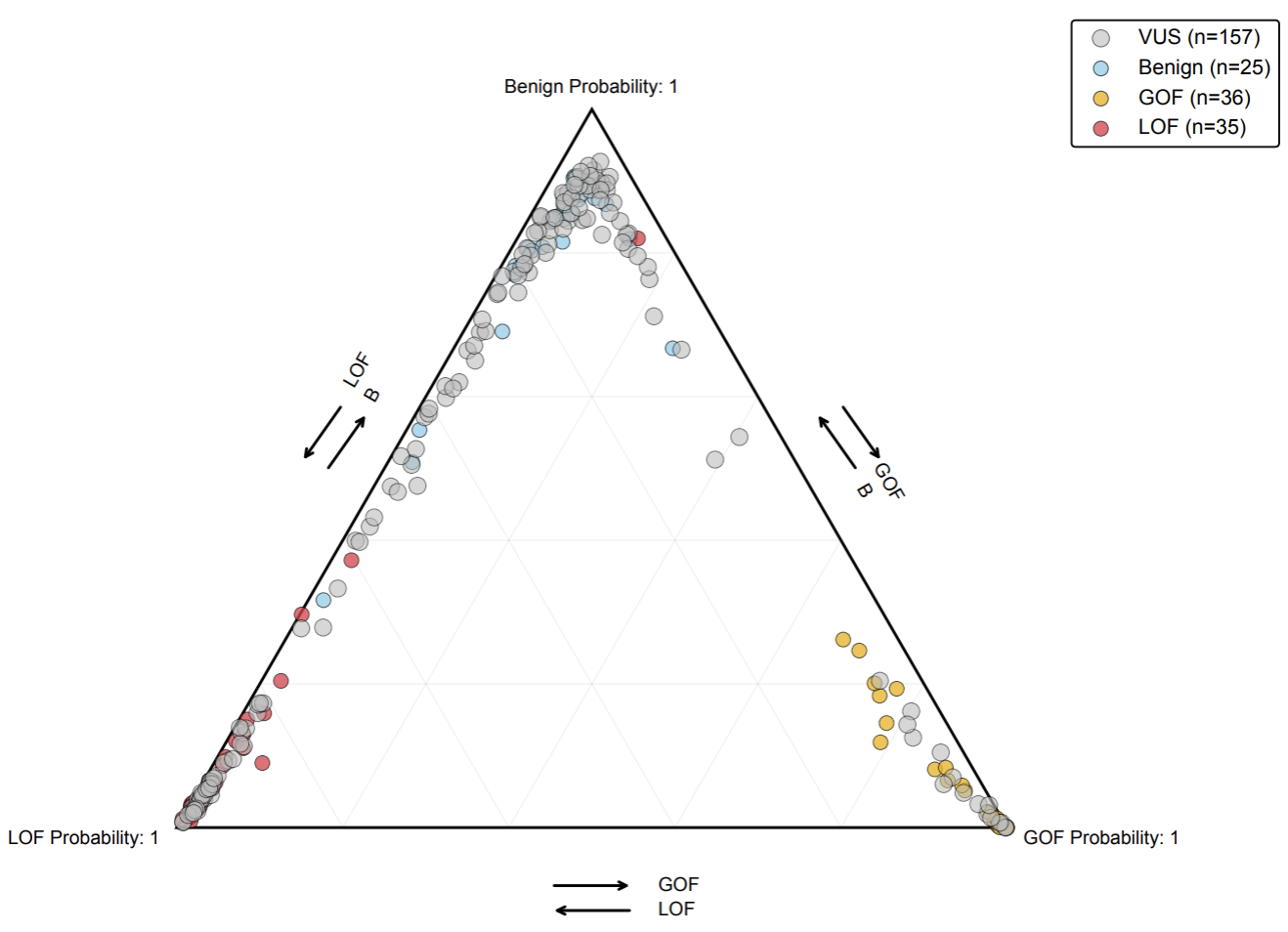

Legend: Figure S6: Predicted class probabilities for annotated variants and VUS. Ternary plot of predicted class probabilities for n=253 variants (dots), positioned by p(Benign), p(GOF) and p(LOF), with proximity to a vertex indicating confident assignment: gray, VUS (n=157); light blue, benign (n=25); gold, GOF (n=36); red, LOF (n=35). Arrows indicate the direction of increasing probability along each axis. As in Figure 3C, with VUS added.

Figure S7

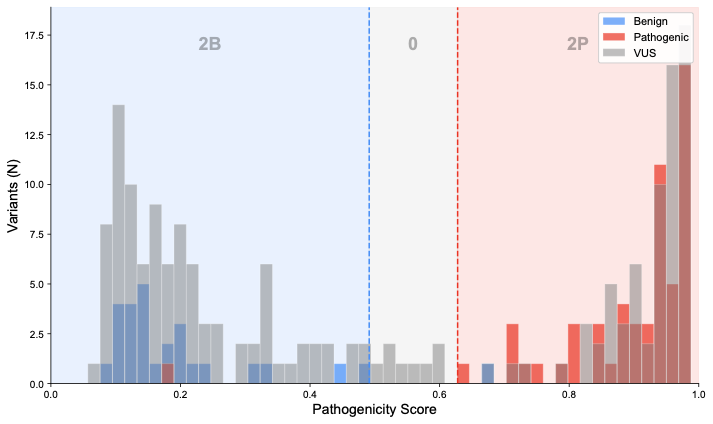

Legend: Figure S7: Calibration of mechanism predictions for Sherloc evidence assignment. Distribution of pathogenicity scores for CASR variants, shown as overlapping histograms by clinical classification: blue, benign; red, pathogenic; gray, VUS. Dashed lines, score thresholds at which the classifier reaches 95% NPV (blue) and 95% PPV (red); shaded regions indicate the corresponding Sherloc evidence assigned to variants in each range (2B, two benign points; 0, no points; 2P, two pathogenic points). No variant fell in the 80% PPV/NPV range corresponding to one point.
